# Cross-Species Comparison of Topologically Associating Domains (TADs) in Cereals Reveals Their Role in Genome Stability During Evolution

**DOI:** 10.64898/2026.08.13.744466

**Authors:** En Li, Liangliang Huang, Junpeng Shi, Guanghui Xu, Hongbing Liu, Weiwei Jin, Yongqiang Wang, Sha Tang, Xianmin Diao, Weibin Song, Beibei Xin, Jinsheng Lai, Jian Chen

**Author notes:** These authors contributed equally to this work (En Li, Liangliang Huang).

## Abstract

Topologically associating domains (TADs) are essential structural and functional modules of the genome that play a crucial role in regulating gene expression. In this study, we systematically investigated the conservation and evolution of TADs in five closely related crops, including maize, sorghum, coix, foxtail millet and broomcorn millet. Our results show that 74% of TAD boundaries are conserved between two inbred maize lines, B73 and Mo17, and that approximately 50% or more of TAD boundaries are conserved across different crop species. TAD number remains relatively stable in the face of changes in genome size. However, the length of TADs varies depending on genome size. Furthermore, we found that large-scale transposable element expansion leads to TAD expansion, while chromosomal inversions lead to TAD fusion and the formation of new TAD boundaries. Frequent chromatin interactions between subgenome chromosomes occur after whole-genome duplication. Moreover, we also found that crossovers are enriched at TAD boundaries in maize, indicating the importance of TADs as a fundamental unit during species evolution. Overall, our study provides insights into the conservation and evolution of TADs in crop genomes and their roles in genome organization and function.

## Introduction

The chromatin of eukaryotic in the nucleus is folded into a highly organized three-dimensional (3D) structure closely related to functional DNA-dependent processes such as DNA replication and transcription [1-5]. In the past decade, chromosome conformation capture methods have revealed key features of genomic organization, particularly the high-throughput version known as Hi-C, which allows the identification of chromatin contacts on a genome-wide scale [6]. Hi-C reveals the general principles of chromosome folding and is able to identify genomic domains with hundreds of kilo-bases and increased self-interaction frequency, known as topological association domains (TADs) [7, 8]. Loci within TADs contact each other more frequently, and TAD boundaries act as insulators to prevent the interactions of loci in different TADs [7, 9]. In mammals, TADs have been reported to restrict the interaction of cell-type specific enhancers with their target genes [2]. Some studies have found that the disruption of TADs can cause ectopic regulation of important developmental genes, leading to genetic diseases [9].

Interestingly, TADs are stable across mammalian cell types and differentiation [7, 10, 11]. TADs are not only found in mammalian genomes, but similar domain organization has been identified in the genomes of non-mammalian species, such as Drosophila [12], zebrafish [13], Caenorhabditis elegans [11], and yeast [14]. This inspired the studies to compare the TAD structure across different species. The reported results suggested that TAD boundaries are largely conserved between human and mouse genomes [7]. About 45% of TAD boundaries in mouse cells occur at homologous regions in human cells. Similarly, 54% of TAD boundaries identified in human cells were also called in homologous positions in mouse genomes [7]. Although 3D genomics has been extensively explored in animals, it has started relatively late in plants in recent years [15, 16]. TAD structures have been reported to exist in various plant species with large-size of genomes, including maize, sorghum and foxtail millet [17, 18]. However, the role of TADs in plants has not been reported, which is considered controversial. Whether TADs are conserved in plants has not been examined yet.

The relatively close (<50 million years old) relationships among diverse cereals make the cereals particularly attractive for comparative studies of plant genome evolution [19]. In order to give a comprehensive examination of TAD conservation between species, we use Hi-C data of five genetically related cereal species including coix, sorghum, maize, broomcorn millet and foxtail millet. The comparative analysis of TADs between evolutionary cereal species is a powerful tool for the delineation of evolutionary-conserved features of chromatin organization and the exploration of TAD functions. We collected the Hi-C data of these species, which enabled us to compare TADs between species. We studied the effects of chromosome inversion, large-scale transposon amplification and whole genome duplication on TADs during the genome evolution, as well as the relationship between crossover and TADs. Our study sheds light on the role of TADs in the evolution of genomes and the maintenance of genomic stability.

## Results

### TADs are conserved between different genotypes in Maize

To examine whether the topologically associating domains (TADs) are conserved in inbred lines with different genotypes, we constructed a Hi-C library with more than three billion read pairs in the endosperm of Mo17. The Hi-C map of Mo17 showed similar patterns as that of B73 (**Fig. 1a**) [17]. Strong interaction signals were displayed along the main diagonal. The Hi-C map presented as ‘X’ shape **(Fig. 1a)**, implying that the chromosome arms from the same and different chromosomes interacted with each other. The identification of TADs was conducted by HiTAD software [20] at 40 kb resolution (**Supplemental Fig. 1, and 2**), which can get the hierarchical structure of TADs (**Fig. 1b**). If we only consider the smallest TADs without overlap, there are 4680 TADs in Mo17 with a median length of 400 kb (**Supplemental Fig. 4a, 4b**).

**Figure 1.**
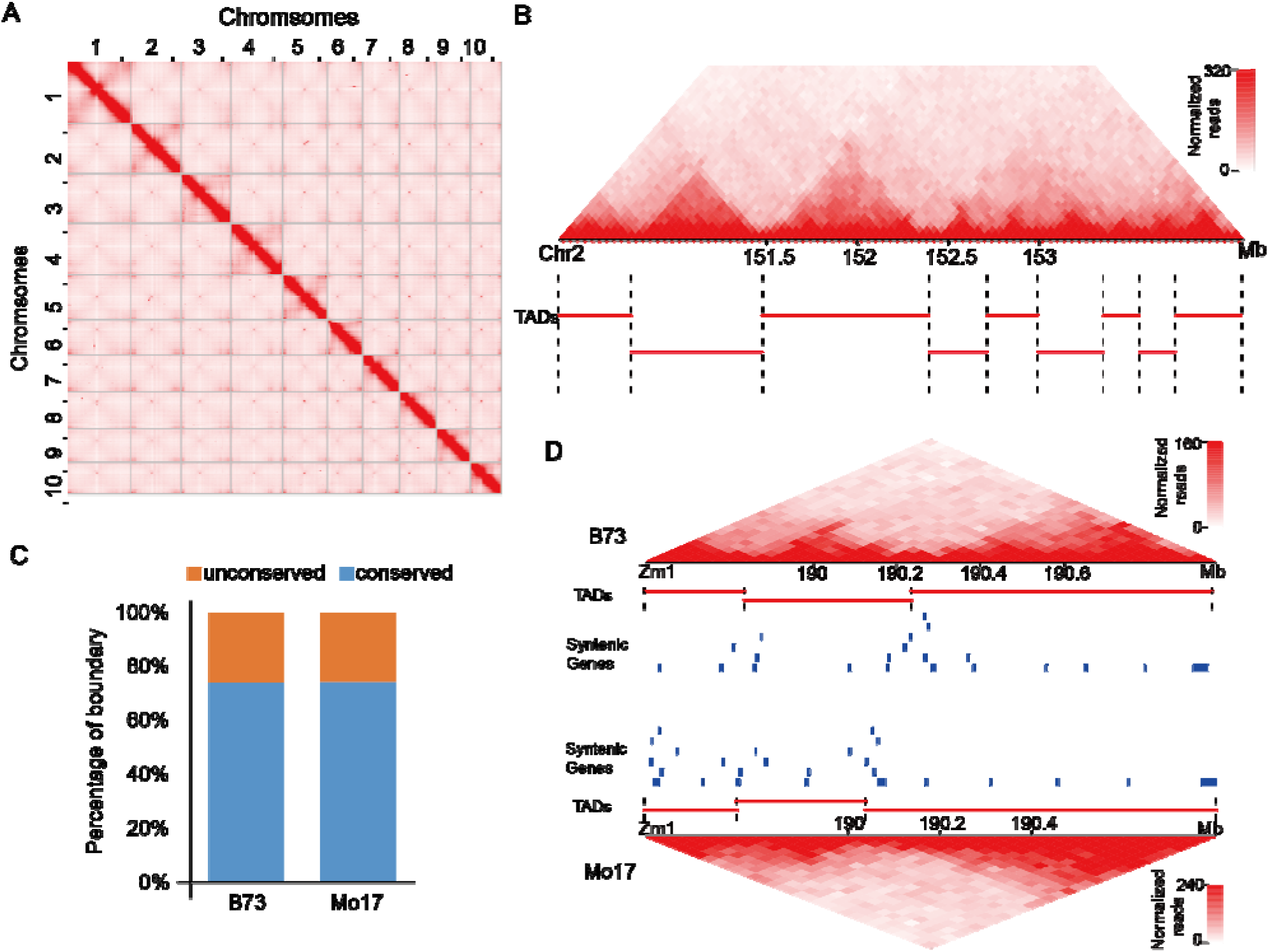
Identification of conserved TAD boundaries and TADs between two maize inbred lines, B73 and Mo17. (A) The genome-wide interaction map of Mo17. (B) Example of TADs in Mo17. The interaction frequency of a region is shown on top. TADs are indicated by red lines below. (C) Percentages of conserved and non-conserved TAD boundaries between B73 and Mo17. (D) Example of three contiguous conserved TADs on a syntenic region in B73 (upper-part) and Mo17 (lower-part). TADs and syntenic genes are indicated by red lines and blue rectangles, respectively.

Previous study in maize have shown that there are about 79%-86% TAD boundaries are conserved between the leaf and endosperm of B73[18]. To further study the conservation of TADs between different genotypes of maize, we identified the TADs by the same methods using the published Hi-C data in the endosperm of B73 [18]. TAD boundary regions with at least one syntenic gene pair between B73 and Mo17 were identified as conserved. We defined that TAD boundaries with syntenic genes in each genotype of maize were analyzable ones. There are 81% and 82% boundary regions that were analyzable in B73 and Mo17, respectively, among which about 74% are conserved in both B73 and Mo17 (compared to 21.0% and 29.0% at random, *p*-value <2.2×10^−16^, Fisher’s Exact Test) (**Fig. 1c**). Our results suggested that the majority of TAD boundary regions were also shared between different genotypes of maize. Next, we studied the conservation of TADs between B73 and Mo17 based on the conservation of boundary, and classified TADs into three groups including completely conserved TADs (CCDs), incompletely conserved TADs (ICDs) and non-conserved TADs (NCDs). The CCDs had two conserved boundaries, while the ICDs had only one conserved boundary. And the NCDs were without conserved boundaries. There were 24% CCDs and 80% TADs with at least one conserved boundary in both B73 and Mo17 (**Supplemental Fig. 4c**). The lengths of conserved TADs between B73 and Mo17 were similar (**Supplemental Fig. 4d**). These results suggested that TADs were conserved not only between different tissues, but also between different inbred lines in miaze.

### TADs are conserved between different species of cereals

Studies in mammals have shown that TADs are conserved in different species [7, 21]. To explore the conservation of TADs in the evolution of plants, we collected Hi-C data of five related cereals species with divergent less than ∼25 Mya, including maize[17], sorghum[17], coix[22], foxtail millet[17], broomcorn millet [23] (**Supplemental Fig. 5**). Then, TAD conservation analysis was conducted between closely related species (sorghum *vs.* coix, foxtail millet *vs.* broomcorn millet and sorghum *vs.* maize) based on syntenic regions, which effectively reduced the noise of comparative TAD analysis among different species. Using the same methods mentioned in comparing TADs between B73 and Mo17 in maize, we identified conserved TAD boundaries and TADs between species, and found that most TAD boundaries are shared between species (**Fig. 2a**). About 66% and 63% of TAD boundary regions were analyzable in sorghum and coix, respectively, **among which 52% and 53% were conserved**. And about 50% of TAD in both sorghum and coix had at least one conserved boundary, among which about 10% of TADs were completely conserved between sorghum and coix. As for the comparative TAD analysis between foxtail millet and broomcorn millet, about 85% and 75% of TAD boundary regions were analyzable, among **which 70% and 47% were conserved**, respectively. And about 77% and 50% of TADs are conserved in foxtail millet and broomcorn millet, respectively, among which 18% and 8% were completely conserved. For the comparative TADs analysis between sorghum and maize, the results showed that about 51% and 70% of TADs boundary were analyzable in sorghum and maize, respectively, **among which 58% and 47% were conserved.** And around 43% and 41% TADs had at least one conserved boundary in sorghum and maize, respectively, among which 6% were completely conserved in both species (**Fig. 2a, 2b, Supplemental Fig. 6**). What’s more, the number of TADs stayed stable on the condition of stable genome ploidy in evolution (**Supplemental table. 1**). Together, our analysis supported that TADs were conserved in the evolution of cereals.

**Figure 2.**
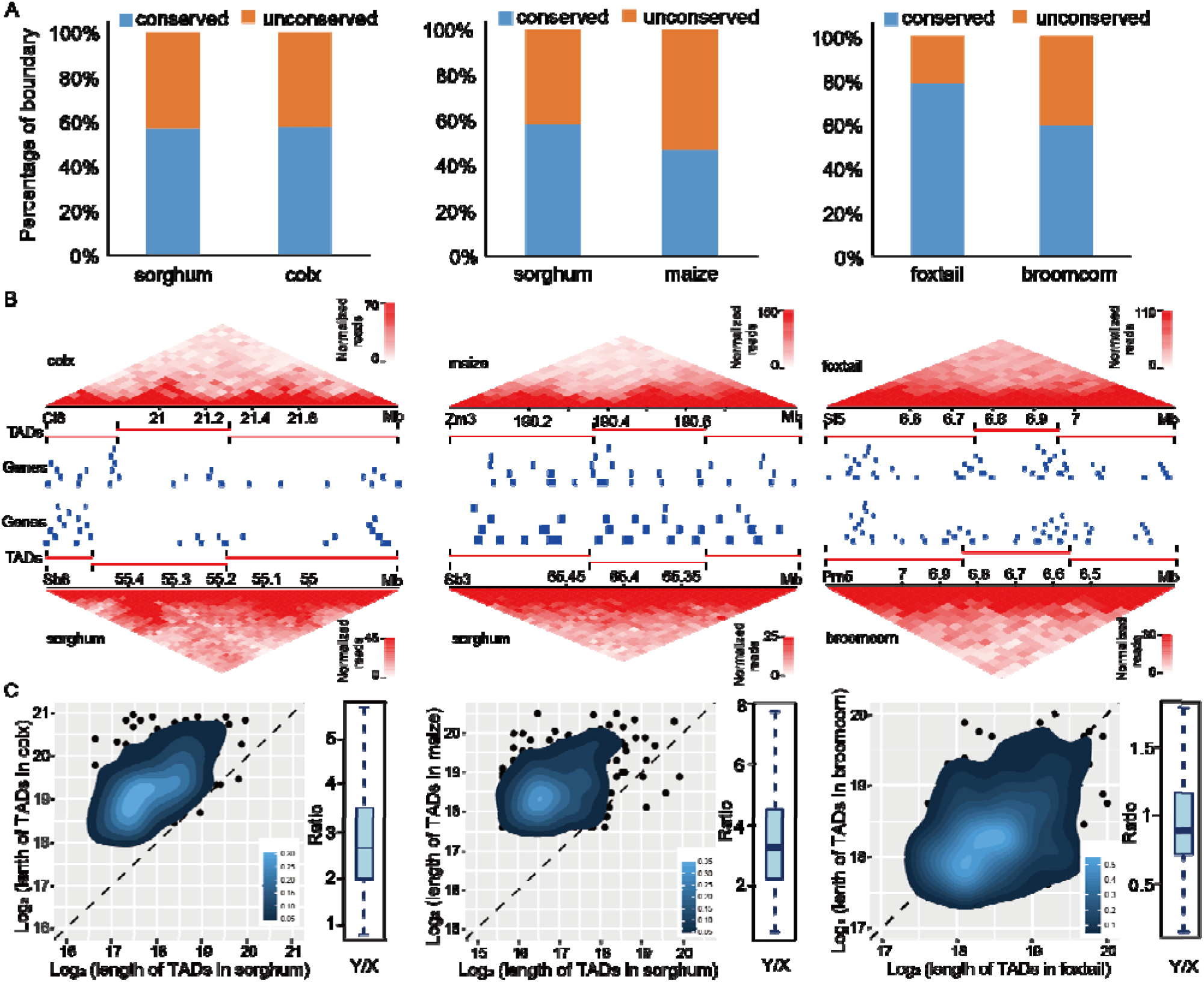
Identification of conserved TAD boundaries and TADs between cereals species. (A) The three bar plots from left to right showed the proportion of conserved and non-conserved TAD boundaries between sorghum and coix, sorghum and maize, foxtail millet and broomcorn millet, respectively. (B) Examples of conserved TADs on a syntenic region between sorghum and coix, sorghum and maize, foxtail millet and broomcorn millet (from left to right panel), respectively. TADs and syntenic genes are indicated by red lines and blue rectangles, respectively. (C) The heatmaps from left to right showed the lengths comparison of conserved TADs between sorghum and coix, sorghum and maize, foxtail millet and broomcorn millet, respectively. The box plot on the right of heat map showed the ratio of conserved TAD length between corresponding two species.

To further characterize the conserved TADs in evolution, we compared the length of conserved TAD pairs between species. The length of TADs in coix was much longer than the conserved ones in sorghum, and the median value of the length ratio between coix and sorghum was 2.67 (**Fig. 2c**). This change may be a result of the drastic genome expansion of coix genome mainly deals to the activity of transposable elements, compared to sorghum [22]. While the lengths of conserved TADs between foxtail millet and broomcorn millet were similar (**Fig. 2c**). The main change of the broomcorn millet genome relative to the foxtail millet genome is that its genome has undergone a whole genome duplication (WGD) event, without strong transposon expansion and chromosome fusions, which makes the chromosome number and genome size of the broomcorn millet genome twice that of the foxtail millet genome [23]. That may cause the change of TAD number, but not the change of TAD length between broomcorn millet and foxtail millet. So the number of TADs identified in broomcorn millet was nearly twice that in foxtail millet, and the length of TADs was almost unchanged between them (**Supplemental table. 1**). As for the comparison of TAD length between sorghum and maize, the results showed that the length of TADs in maize was always much longer than that in sorghum, and the median value of length ratio between maize and sorghum was 2.47 (**Fig. 2c**). Comparative genomic analysis has revealed that maize is an ancient tetraploid and experienced strong chromosome fusions, gene loss and transposon expansion [24, 25]. The change of genome may lead to the change of TAD length and number between maize and sorghum. Taken together, our results indicated that TADs were conserved in different species, but some genome events can bring the change of TAD length in the evolution. Since the TADs are conserved in plants, we next want to study the events that may impact the integrity or the structure of TADs, including transposon expansion, chromosome inversion and crossover.

### Large-scale Transposon expansions cause the expansion of TAD length

To study the effect of transposon expansion on TAD conservation in evolution, we focused on chromosome 3 of coix (cl3) and its corresponding chromosome 10 of sorghum (sb10). On the comparative map of homologous sequences between sorghum and coix genome produced by Mummer software [26], it can be observed that the cl3-sb10 mainly consisted of two segments with distinctly different slopes (**Fig. 3a**). The slope corresponding to cl3:1-109073628 (named as strongly expansive region, SER) was smaller than that corresponding to cl3:109073629-179537263 (named as weakly expansive region, WER) (**Fig. 3a**), indicating that the genome expansion in the former segment was greater than that in the latter [22]. Through exploring the conservation of TADs on the two segments of cl3, we found that the TAD conservation of SER was slightly smaller than that of WER (**Fig. 3b, Supplemental Fig. 9a**). For SER, nearly 74% and 49% of TAD boundary regions were analyzable in sorghum and coix, respectively, among which 52% and 40% were conserved. And there were 61% and 32% TADs with at least one conserved boundary in sorghum and coix, respectively, among which 8% and 5% were completely conserved. As for WER, about 64% and 60% of TAD boundary regions were analyzable in sorghum and coix, respectively, among which 57% and 50% were conserved. And there were 53% and 52% TADs with at least one conserved boundary in sorghum and coix, respectively, among which 9% and 8% were completely conserved (**Fig. 4b, Supplemental Fig. 9a**). To further validate our hypothesis that the genome expansion can cause the expansion of TADs, we compared the length ratios of TADs between coix and sorghum on SER and WER. The result showed that the ratio of SER was significantly larger than that of WER, indicating that the expansion of TADs in SER was remarkably greater than that in WER (**Fig. 4c, 4d**). Moreover, both the boundary regions and inner regions of TADs on SER showed stronger transposon expansion than that on WER (**Supplemental Fig. 9b**). Despite the different transposon expansion levels between SER and WER, we did not find obvious different expression correlations between syntenic gene pairs located on these two segments (**Supplemental Fig. 9c**), which suggested that TADs may play a role in buffering the effects of transposon amplification on gene expression during species evolution.

**Figure 3.**
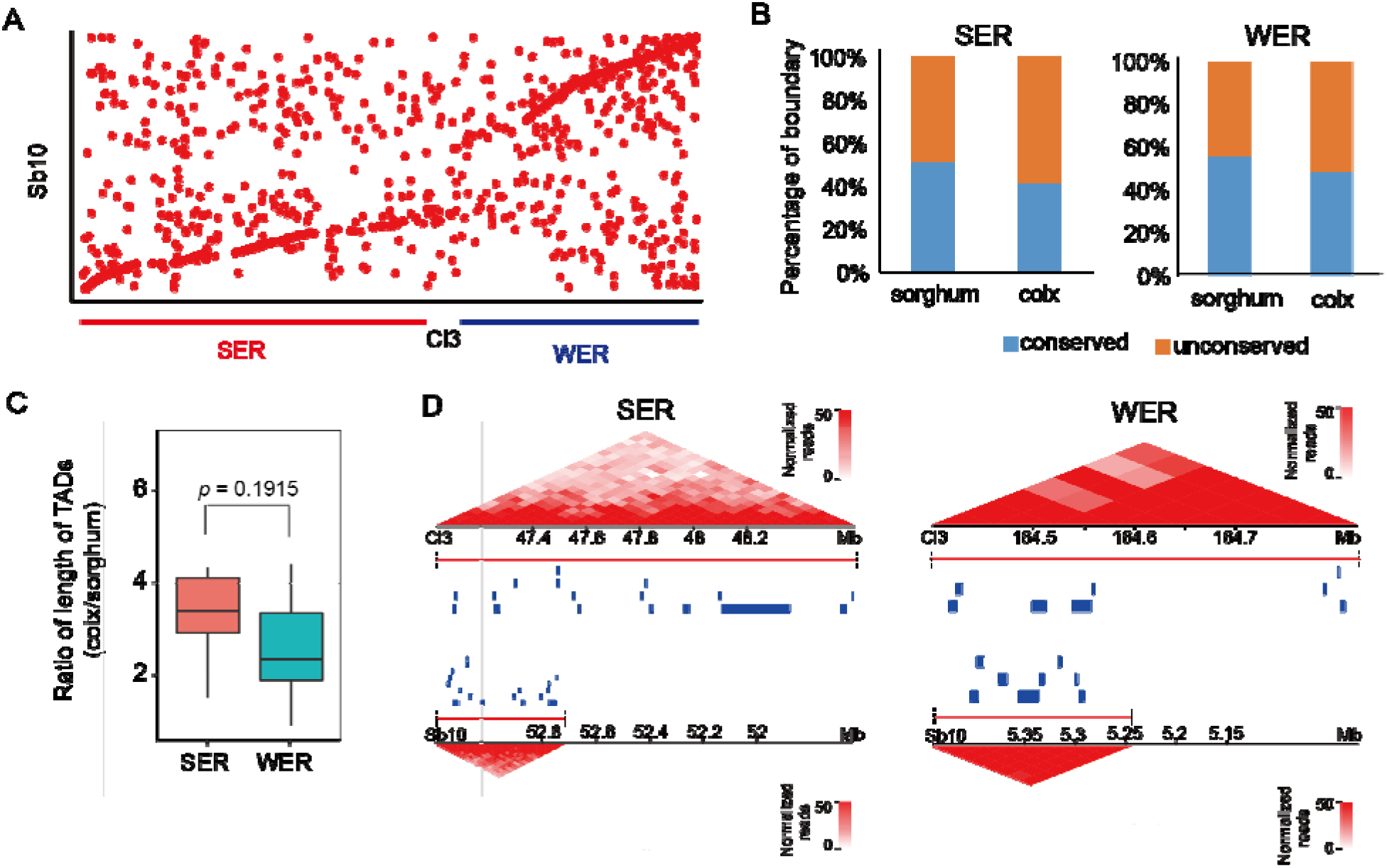
Genome expansion lead to the expansion of TADs in coix. (A) Genome comparison between Coix (cl3) and Sorghum (sb10). Each dot represented a homologous sequence reported by Mummer. (B) The bar plots showed the proportion of conserved and non-conserved TAD boundary in SER and WER between sorghum and coix. (C) Comparison of the ratio of TAD length between coix and sorghum in SER and WER. (D) Examples of changed length of conserved TADs on a syntenic region between sorghum and coix located in SER and WER, respectively. TADs and syntenic genes are indicated by red lines and blue rectangles, respectively.

**Figure 4.**
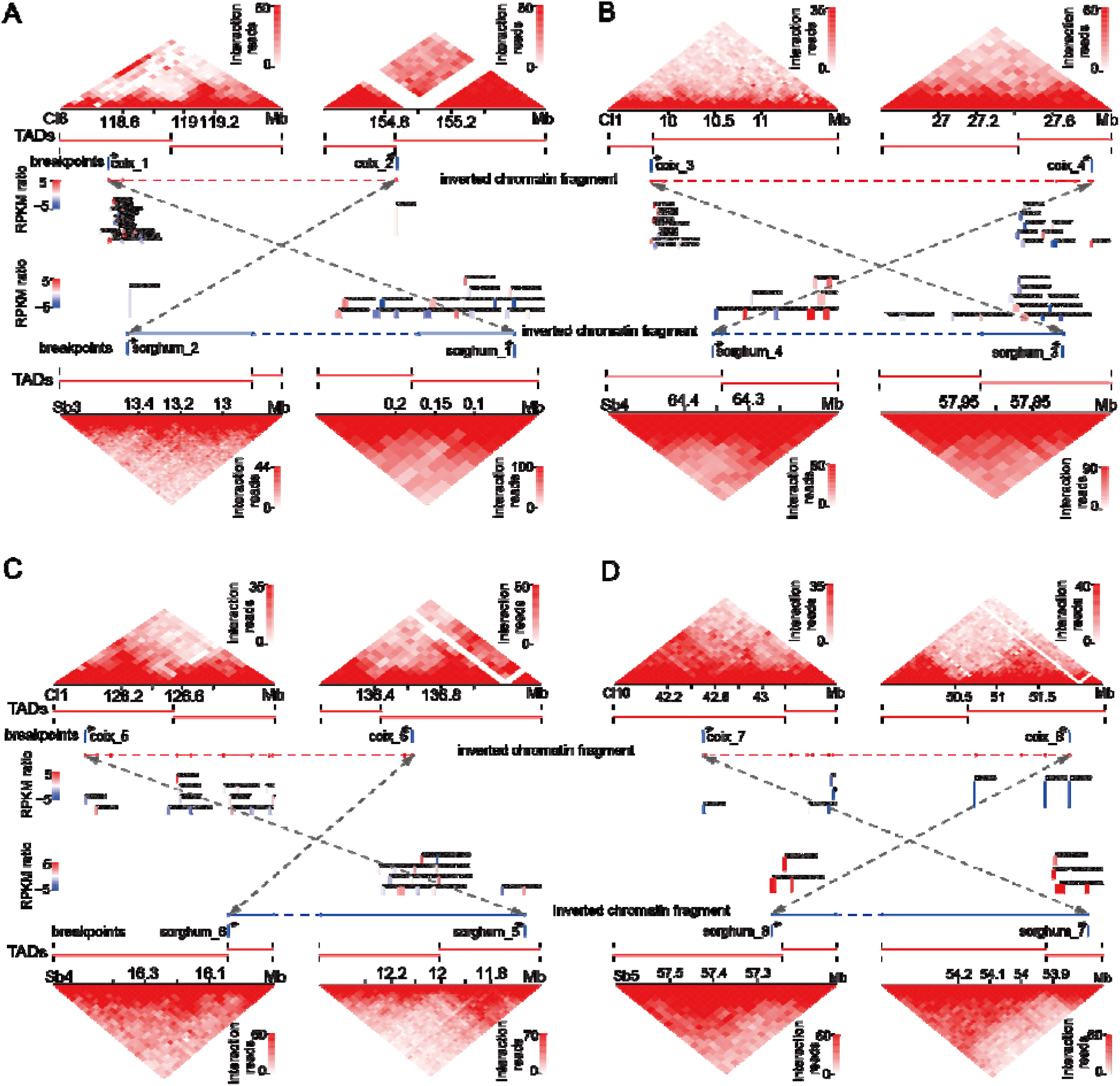
TAD fusion and the formation of TAD boundary at the breakpoints of chromosome inversion between coix and sorghum.

### Large chromatin inversion leads to TAD fusion or the formation of new TAD boundaries

Chromatin inversion is another common event in genome evolution, but its effect on TAD has not been investigated in evolution. In this study, we chose the seven reported chromosomal inversion events occurred between the coix and sorghum [22], to study the effects of large chromatin inversion on the three-dimensional chromatin structure and on the expression of surrounding genes. The breakpoints of these inverted chromosome fragments were mostly (11/14) occurred in inner regions of TADs, among which 64% (7/11) leaded to TAD fusion, and the remain caused the formation of new TAD boundary (**Fig. 5**). The genes surrounding the breakpoint of TAD fusion can be up-regulated or down-regulated due to the possibility of enhancer hijacking or loss (**Supplemental Fig. 10**). As for the new TAD boundary formed at the breakpoint, it was always accompanied by the up-regulated expression of surrounding genes (**Supplemental Fig. 10 C-D**), which was consistent with the reported phenomenon that genes with high expression levels are enriched at the TAD boundary [7]. Besides, TADs within the inverted fragment were still highly conserved between sorghum and coix (**Supplemental Fig. 11**), comparable to the genome-wide conservation level observed between these species (**Fig. 2A**).

**Figure 5.**
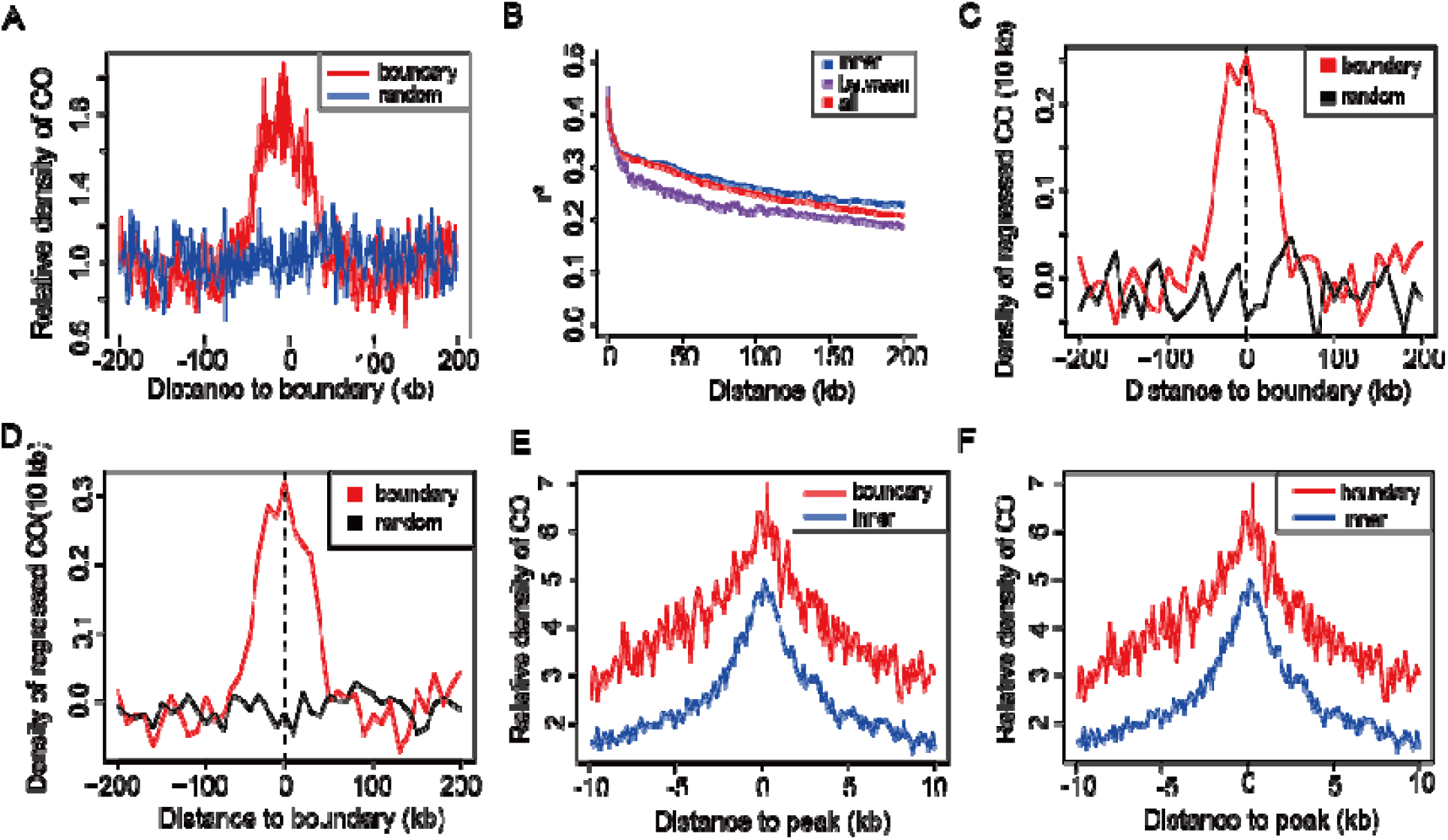
CO show an enrichment at TAD boundary. (A) The relative density of CO around TAD boundary in maize. (B) The LD decay between and w thin TADs in maize. (C) The relative density of the regressed of CO by gene density around TAD boundary in maize. (D) The relative density of the regressed of CO by ATAC peak around TAD boundary in maize. (E) Compare the relative density of CO around gene located in boundary and inner of TADs. (F) Compare the relative density of CO around ATAC peak located in boundary and inner of TADs.

### Crossovers are enriched in the boundary of TADs in maize

Although various features of TAD boundary have been reported in both mammals and plants, enriched housing-keeping genes, highly expressed genes and active epigenetic modifications [7, 17, 27], the biological functions of TADs remain to be explored. To explore the biological function of TADs in genome evolution, we examined the distribution of crossovers (COs) around the TAD boundary to study the contribution of TADs to genomic recombination. We identified the recombination breakpoints in maize, which were defined as crossovers, using a collection of 25 biparental populations that consist of 5,230 recombinant inbred lines derived from the cross of 25 diverse maize accessions with a common parent B73 [28]. Crossovers showed an enrichment in TAD boundary regions in maize (**Fig. 5a**). A high recombination rate can cause rapid Linkage disequilibrium (LD) breakdown [29-31]. We found that LD decayed faster between loci located on different TADs than those located in the same one (**Fig. 5B and Supplemental Fig. 12**), indicating that loci within the same TAD tend to co-segregate and is consistent with the enrichment of crossovers at TAD boundaries. Previous studies have reported that crossovers tend to predominantly happen within open chromatin regions, gene promoters and terminators, and regions with low nucleosome density [32-34]. Genes and ATAC-seq peaks also showed enrichments in the boundary regions of TADs in plants (**Supplemental Fig. 13A and B**). To eliminate or mitigate the effect of genes on boundary regions for crossover, we regressed the crossover frequency by gene coverage on the genome and still observed an enrichment of crossover in the boundary regions (**Fig. 5C**). Further, we compared the distribution of crossover around the gene in boundary regions and inner regions of TADs. Although the crossover showed enrichments on genes in both regions, the frequency of crossover is much higher around genes in boundary regions than in inner regions of TADs (**Fig. 5D**). Leveraging ATAC-seq data, we further performed a regression analysis of crossover frequency based on chromatin accessibility, revealing a persistent enrichment of crossovers in the boundary regions (**Fig. 5E**). Simultaneously, in comparison to the density of crossovers near ATAC-seq peaks distributed within the internal regions of TADs, ATAC-seq peaks located at the boundaries of TADs exhibit a higher abundance of crossovers (**Fig. 5F**). This implied that TAD tended to be a complete unit in homologous recombination.

## Discussion

In this study, we systemtically investigated the TAD conservation among five genetically related cereal species including coix, sorghum, maize, broomcorn millet and foxtail millet. There were 74% TAD boundaries that were conserved between the two maize inbred lines, B73 and Mo17. About 50% or more of TAD boundaries are conserved between different crop species. The number of TADs stayed stable on the condition of stable genome ploidy in evolution. These results suggest that TADs is a conserved unit of genome and may play a crucial role in the evolution of crop genomes. Another interesting finding regarding plant TADs is their hierarchical organization, with sub-TADs nested within higher-level TADs, consistent with observations in mammals [35-37], suggesting that chromatin may manage the genome through stratification. When studying TAD conservation, especially across species, it is important to consider this hierarchical structure. In addition, the resolution of Hi-C data can affect TAD calling and the interpretation of conservation. Therefore, to avoid complications arising from TADs at different hierarchical levels, we recommend focusing on the conservation of TAD boundaries instead.

Although TADs are generally conserved among species, certain major evolutionary events can influence TAD architecture. We found that large-scale transposable element expansion can increase TAD length without significantly disrupting TAD conservation. By confining the effects of transposable element insertion and expansion to limited genomic units, TADs function as effective buffers that mitigate their impact on overall genome organization and maintain relative genome stability, underscoring the benefits of hierarchical genome organization. Consequently, because different degrees of transposable element expansion lead to substantial variation in TAD length, it is necessary to adjust the resolution used for TAD calling according to the extent of genome expansion when assessing TAD conservation between species. In contrast, large chromatin inversions can disrupt TAD organization, creating rearranged TADs that alter gene regulatory patterns and introduce substantial genomic variation.

Meiotic recombination is an important event driving genomic variation across generations. We found that crossovers are enriched at TAD boundary regions in maize, suggesting that TADs tend to act as intact units during homologous recombination, which may help maintain the regulatory stability of genes within TADs. Double-strands chromatin break (DSB) is the precondition of crossover. Previous study in maize has shown that RAD51 protein is required for DSB repair and homologous recombination, and forms foci on chromosomes at the sites of DSB in maize [38, 39]. We further studied the distribution of DSB in the TAD boundary by using the RAD51 ChIP-seq [38]. Only a slightly higher frequency of DSB on the boundary than in the other regions (**Supplemental Fig. 14A**).Similarly, there was no obvious difference between the distribution of DSB of gene and active regions in boundary and inner regions of TADs (**Supplemental Fig. 14B and C**). This contrasts with findings in humans, where DSBs are enriched at high-level TAD boundaries [40]. The discrepancy may reflect the lack of hierarchical categorization in our TAD analysis, highlighting the need for future studies to explore the properties of TADs at different hierarchical levels in plants. Interestingly, we also observed that longer TADs tend to be enriched for repressive marks, whereas smaller TADs are more likely to carry active marks (**Supplemental Fig. 15**), suggesting that TADs of different hierarchical levels may have distinct functional roles in genome regulation.

## Methods

### Plant Material and Hi-C Library Construction

The maize (Z. mays) inbred line Mo17 was cultivated in the field in Beijing, China, specifically to collect endosperm samples 12 days after pollination. The harvested endosperm was fixed using 2% formaldehyde for 15 minutes, followed by the addition of glycine to quench the crosslinking reaction. Subsequently, the supernatant was removed, and the tissue was dried using blotting paper before being ground to isolate the nuclei.

To proceed, we followed the published Hi-C protocol [10, 41]. In summary, DNA was extracted from the isolated nuclei and digested using MboI. The resulting crosslinked DNA fragments were labeled with biotin-14-dCTP at the ends and then re-ligated using the ligation enzyme. The nuclear complexes were subsequently reverse-crosslinked, and the ligated DNA fragments were extracted. These fragments were sonicated to achieve a size range of 100-500 bp and used for constructing Hi-C libraries. The libraries were sequenced on the Illumina HiSeq platform using the PE 150 sequencing strategy.

### Hi-C Data Analysis

In this study, we utilized the maize B73 (RefGen_v4) [42], Mo17 v1.0 [43], sorghum v3.1, coix v1.0 [22], foxtail millet v2.2, broomcorn millet v1.0 [23] as reference genomes. To align the clean Hi-C reads of each species to their respective reference genomes, we employed the HiC-Pro software [44]. For coix, we utilized a 60 kb bin size for Hi-C matrix analysis and TAD calling. In the case of sorghum and maize, a bin size of 20 kb was employed. For foxtail millet and broomcorn millet, a bin size of 40 kb was used for Hi-C matrix analysis and TAD calling. To ensure sufficient sequencing depth at the chosen resolution, we followed a previously reported method [6], which required the top 80% of the bins to possess no fewer than 1,000 *cis*-interaction (**Figure S3, S5**). The HiTAD software [45] was employed to identify TADs in each species at the corresponding resolution. All Hi-C contact maps were generated using Juicer [46], HiCPlotter software [47], or the Sushi package in R.

### Identification of conserved TADs boundary and TADs between species

We defined the boundary region of TADs identified by HiTAD as the left and right bins adjacent to the TAD boundaries. Throughout the manuscript, the term “boundary” refers to the region encompassing the left and right bins adjacent to the TAD boundaries. The criterion for identifying conserved boundary regions across different species was the presence of syntenic gene pairs. The identifications of syntenygene pairs between coix and sorghum, maize and sorghum, maize and coix, foxtail millet and broomcorn millet were conducted using the CoGe pipeline (https://genomevolution.org/).

Conserved TAD identification is based on the conservation of TAD boundaries. If both boundaries of a TAD are conserved, we define it as a completely conserved TAD. If only one boundary of a TAD is conserved, we define it as an incompletely conserved TAD. If neither of the boundaries of a TAD is conserved, we define it as a non-conserved TAD.

### Transcriptome, Histone Modification, Methylome, and ATAC-seq Analysis

The mapping of RNA-seq reads to the respective reference genome was performed using TopHat [48]. Subsequently, Cufflinks software [49] was utilized to determine the gene expression levels. The calculation of Shannon entropy for each gene in maize was performed as previously described [50]. In order to determine the co-expression level of paralogous gene pairs between the two subgenomes of broomcorn millet, we analyzed the expression patterns of gene pairs using the Pearson’s correlation coefficient in a collected RNA-seq dataset. The dataset comprised a total of 23 diverse samples from broomcorn millet, which was previously reported by Shi et al. [41]. The gene expression data for sorghum and coix were obtained from Liu et al. [22]. BS-seq data were processed using Bismark software [51]. We mapped the ChIP-seq and ATAC-seq data using bowtie2 [52] with default parameters. Reads were first trimmed to 50 bp and only mapped reads with MAPQ ≥ 10 were used for further analysis. PCR duplicates were removed by samtools. For histone modification data, we used MACS2 [53] for peak calling with parameters “--nomodel --broad --broad-cutoff 0.01”. For ATAC-seq data, MACS2 was used with parameters “--nomodel --extsize 70 -q 0.01”.

### Maize chromosome recombination map

To identify recombination breakpoints in maize, we used a collection of 25 biparental populations that consist of 5,230 recombinant inbred lines derived from the cross of 25 diverse maize accessions with a common parent B73. This population, as well as their parental lines, had been genotyped with 12,659,487 SNP markers [28]. Recombination breakpoints were determined as the midpoint positions of two SNP makers that displayed different genotypes (AA, AB, or BB). Analysis of the 25 biparental populations revealed a total of 117,817 unique recombination breakpoints.

### LD analysis

LD measurement parameter R^2^ was used to estimate LD between SNPs with MAF > 0.05 on each chromosome for the entire germplasm set. To see how much the subpopulation would affect the extent of LD, R^2^ was also calculated separately for the tropical and temperate germplasm sets. TASSEL 3.0 software [54] was used to calculate the extent of LD (R^2^) between SNP pairs at *P* = 0.01 in the entire set and each germplasm set.

## Data availability

The Hi-C data for maize Mo17 is accessible via accession number <u>GSE304159</u>. Other data used in the manuscript is publicly available.

## Author Contributions

E.L., J.C., and J.L. designed the research; J.C., H.Z., W.S., B.X., and J.L. collected the plant materials, performed the experiments, and generated the sequencing data; E.L. and L.H. analyzed the Hi-C data; G.X., J.S., H.L., and W.J. participated in the data analysis. E.L. wrote the manuscript. E.L, L.H, J.C. and J.L revised the manuscript.

## Acknowledgments

This work was supported by the National Natural Science Foundation of China [32572347], and the 2115 Talent Development Program of China Agricultural University.

---

**Supplemental Figure 1.**
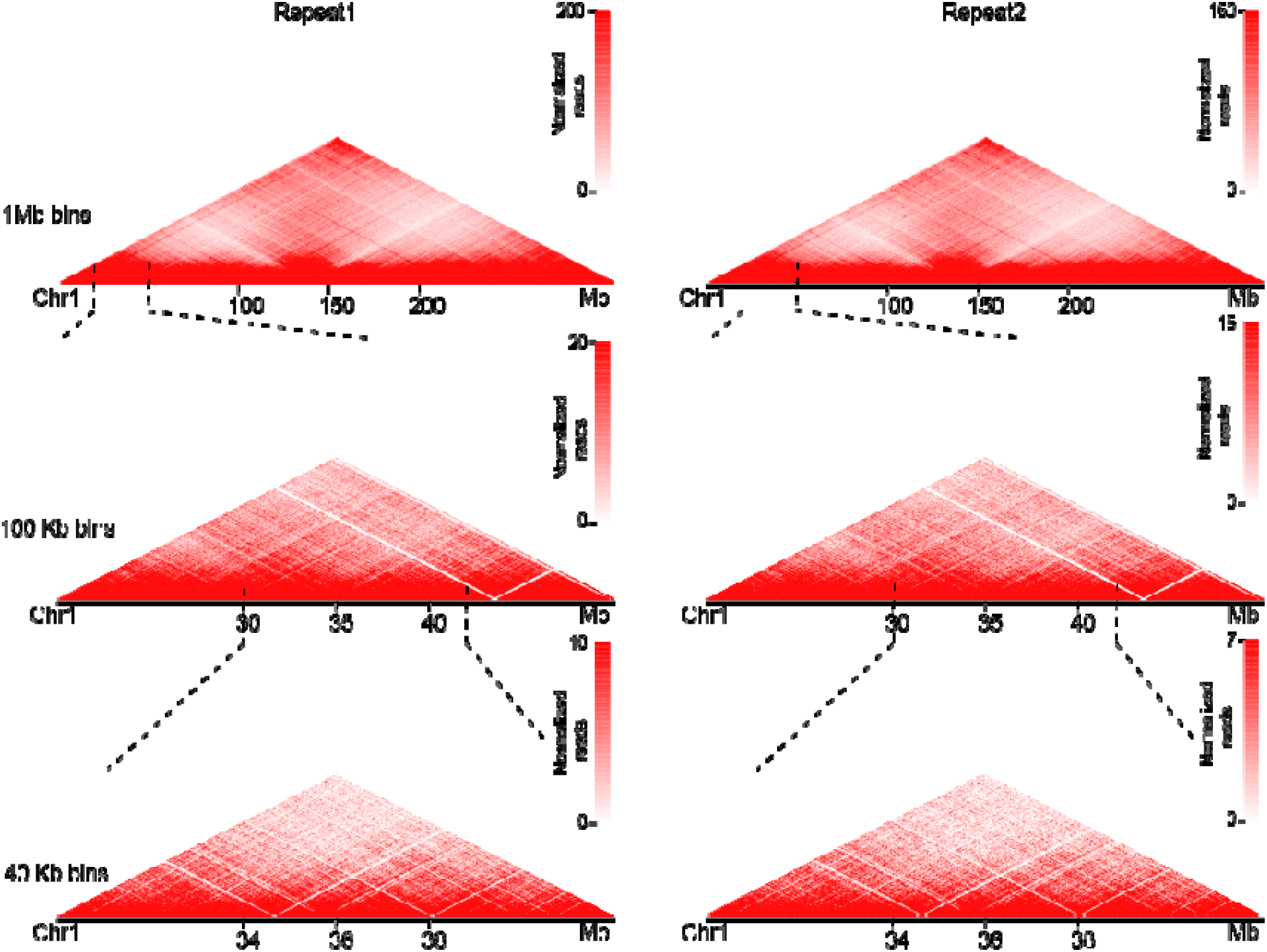
Comparision of Hi-C interaction heat maps between replicates in Mo17, related to Fig1. Contact matrices from chromosome 1 in endosperm of Mo17: the whole chromosome, at 1 Mb resolution (top); 20–50 Mb/100Kb resolution (middle); 30–42 Mb/40 Kb resolution (bottom). Replicate1; Right: Replicate2.

**Supplemental Figure 2.**
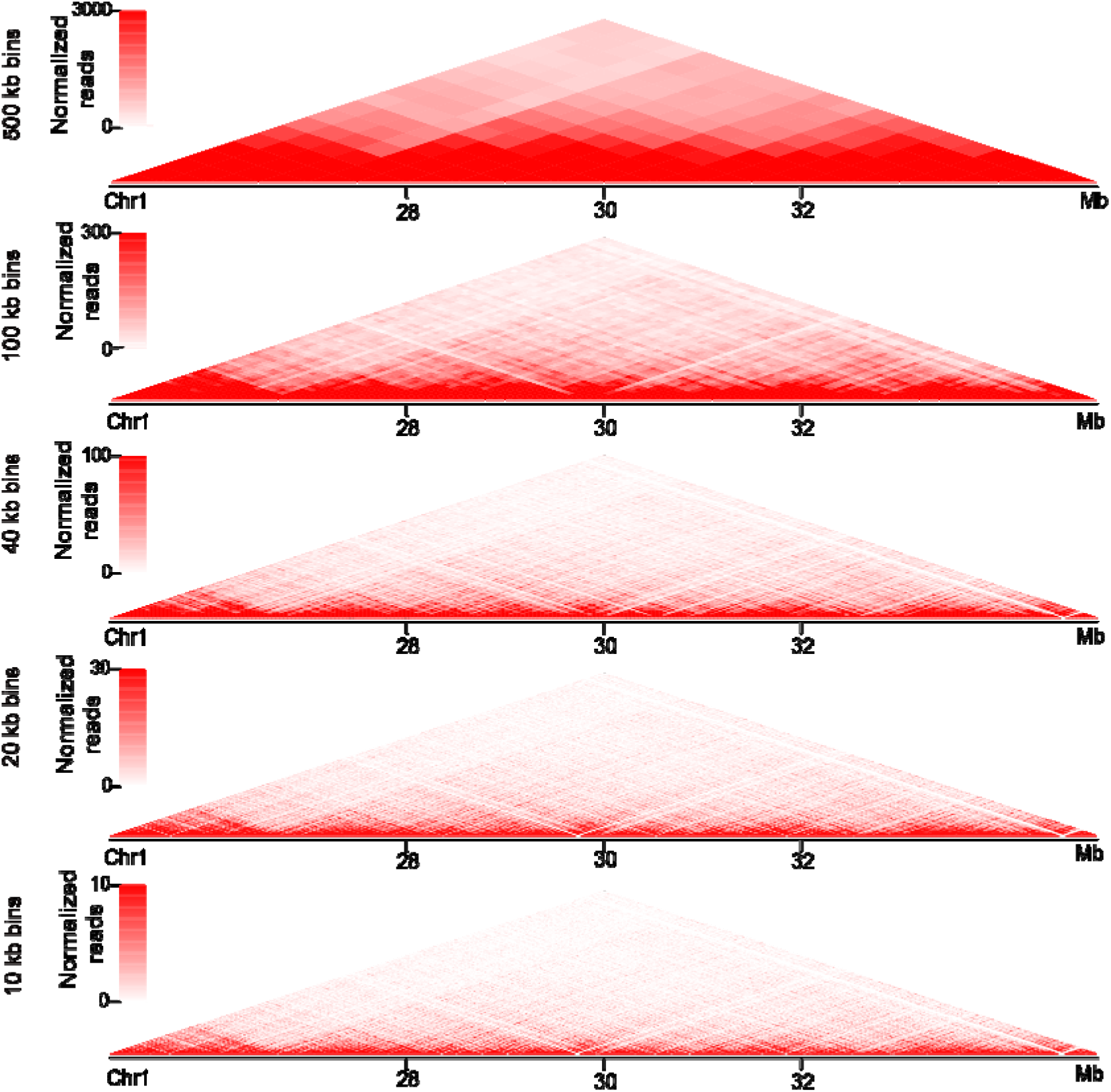
Hi-C interaction heatmaps at varying bin sizes, related to Fig1. Hi-C interaction frequencies are displayed as 2D heatmaps using differing bin sizes over a single locus (25–35 Mb) on chromosome 1.

**Supplemental Figure 3.**
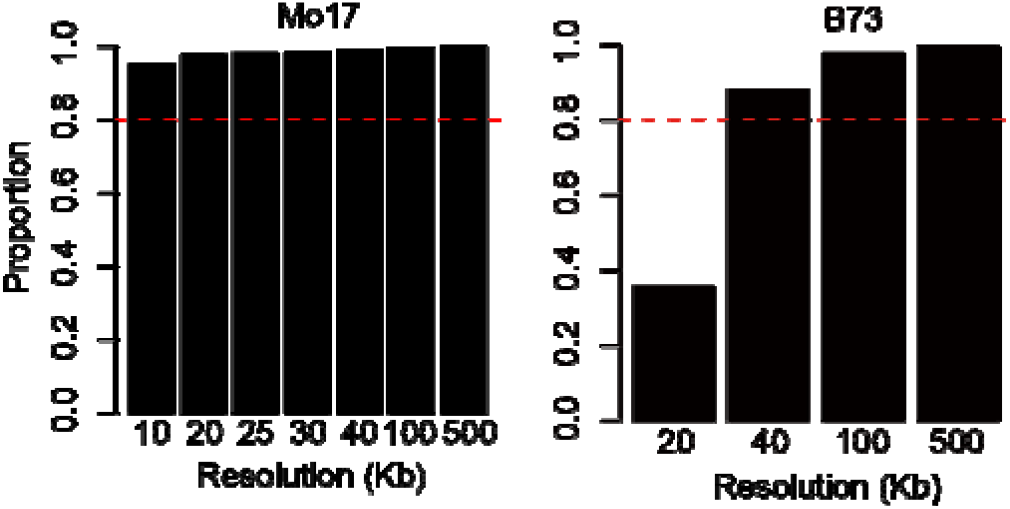
The resolution of Hi-C datas in endosperm of two maize inbred lines, B73 and Mo17, related to Fig1. According to the defined Hi-C resolution [55], the resolution of Hi-C data ensured that 80% of loci have at least 1000 contacts.

**Supplemental Figure 4.**
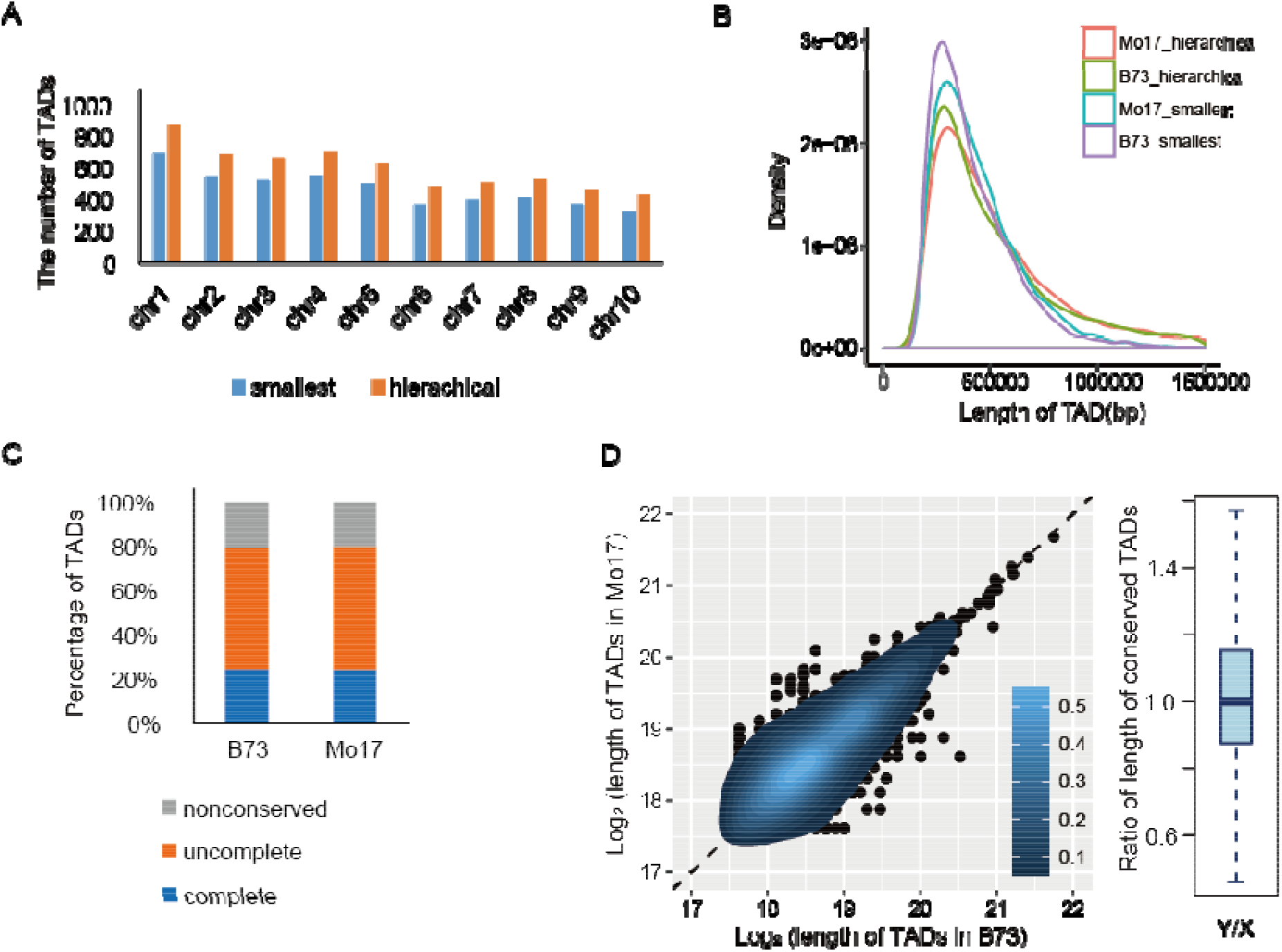
Identification of conserved TADs between two maize inbred lines, B73 and Mo17, related to Fig1. **(A)** Summary of the number of TADs identified in each chromosome of Mo17. **(B)** The length distribution of TADs in B73 and Mo17. **(C)** The bar plots showed the proportion of non-conserved, incompletely conserved and completely conserved TADs between B73 and Mo17. **(D)** The heatmap showed the lengths comparison of conserved TADs between B73 and Mo17. The box plot on the right of heatmap showed the ratio of conserved TAD length between B73 and Mo17.

**Supplemental Figure 5.**
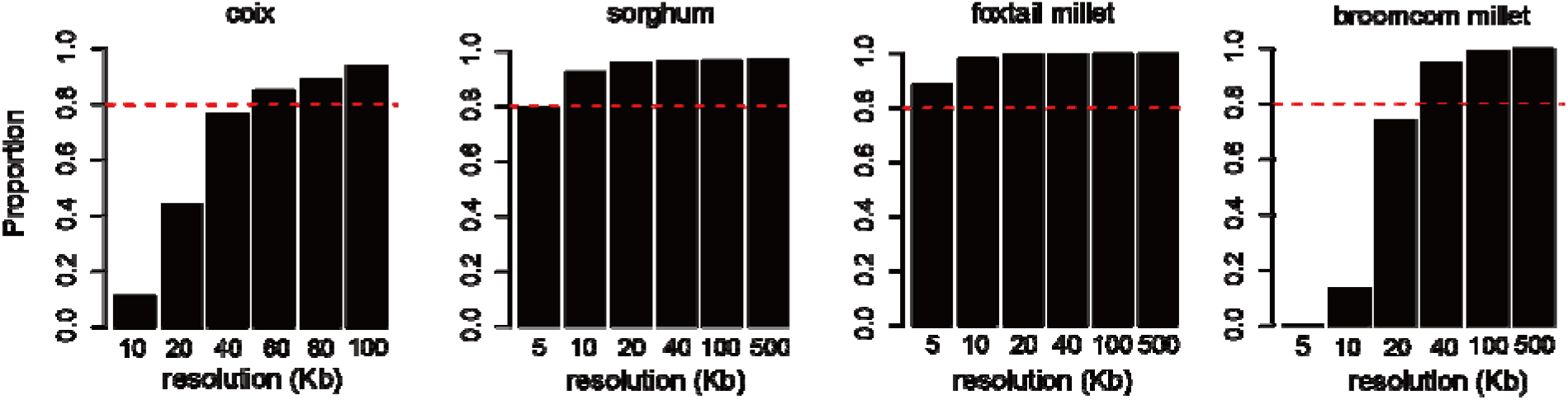
The resolution of Hi-C datas in shoot of each cereals species, related to Fig2. According to the defined Hi-C resolution [55], the resolution of Hi-C data ensured that 80% of loci have at least 1000 contacts.

**Supplemental Figure 6.**
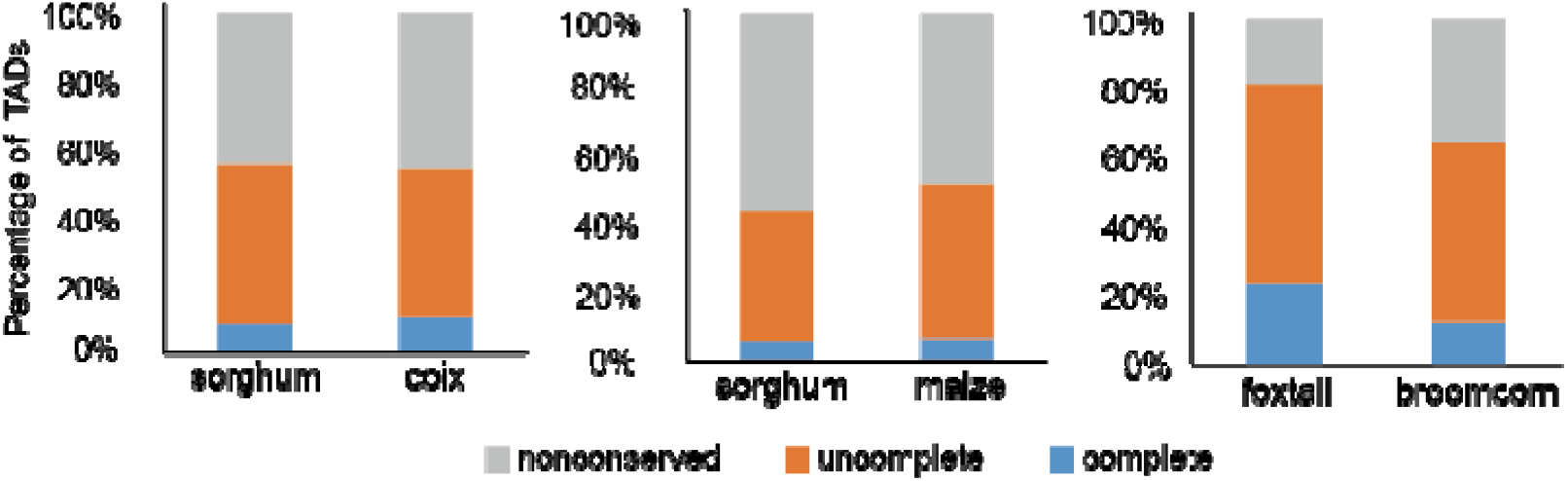
Identification of conserved TADs between cereals species, related to Fig2. The barplots from left to right showed the proportion of non-conserved, incompletely conserved and completely conserved TADs between sorghum and coix, sorghum and maize, foxtail millet broomcorn millet, respectively.

**Supplemental Figure 7.**
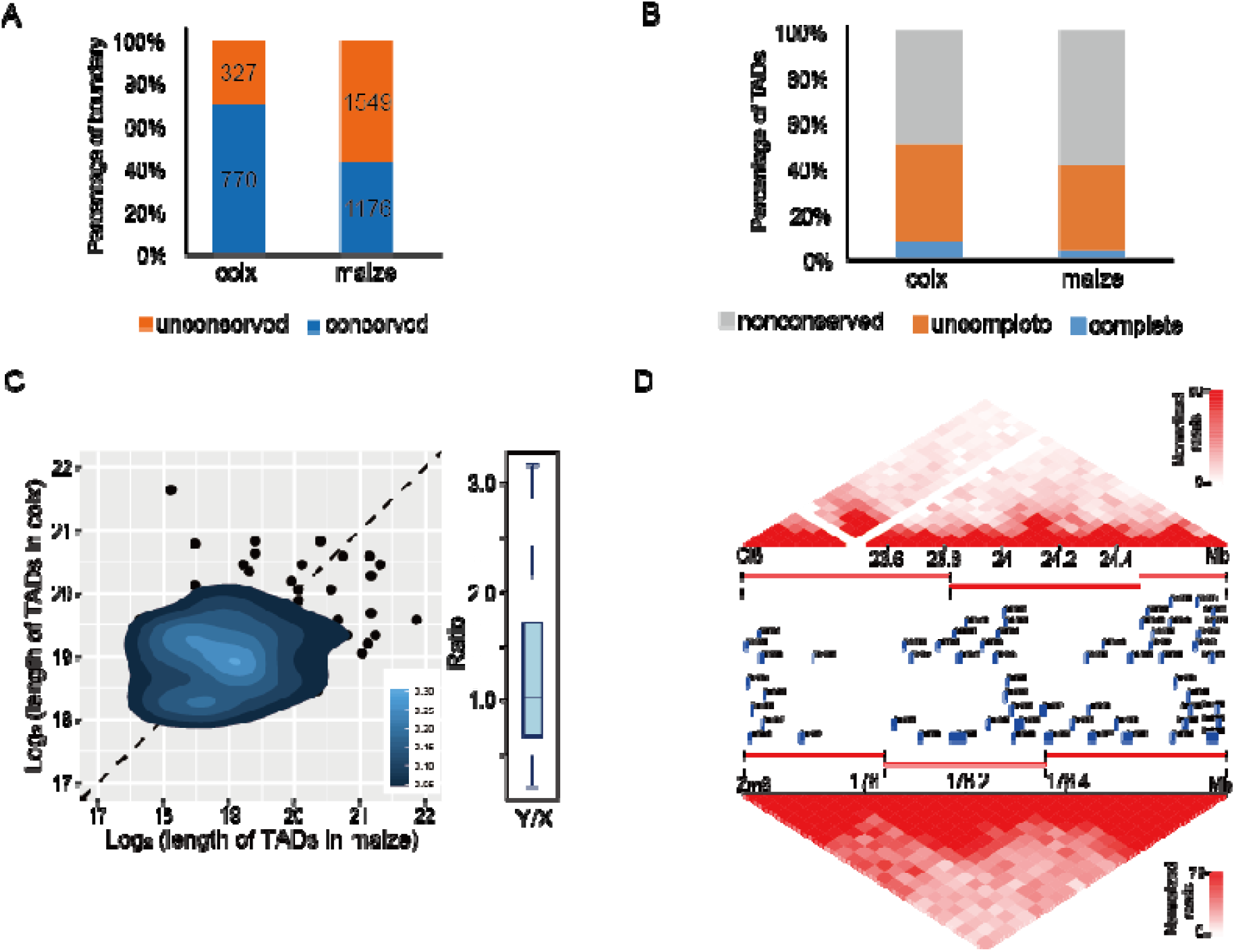
Identification of conserved TADs between coix and maize, related to Fig2. **(A)** Percentages of conserved and non-conserved TAD boundaries between coix and maize. Numbers on the bars indicated the absolute number of TAD boundaries. **(B)** The bar plot showed the proportion of non-conserved, incompletely conserved and completely conserved TADs between coix and maize. **(C)** The heatmap showed the lengths comparison of conserved TADs between coix and maize. The box plot on the right of heat map showed the ratio of conserved TAD length between corresponding two species. **(D)** Example of conserved TADs on a syntenic region between coix and maize. TADs and syntenic genes are indicated by red lines and blue rectangles, respectively.

**Supplemental Figure 8.**
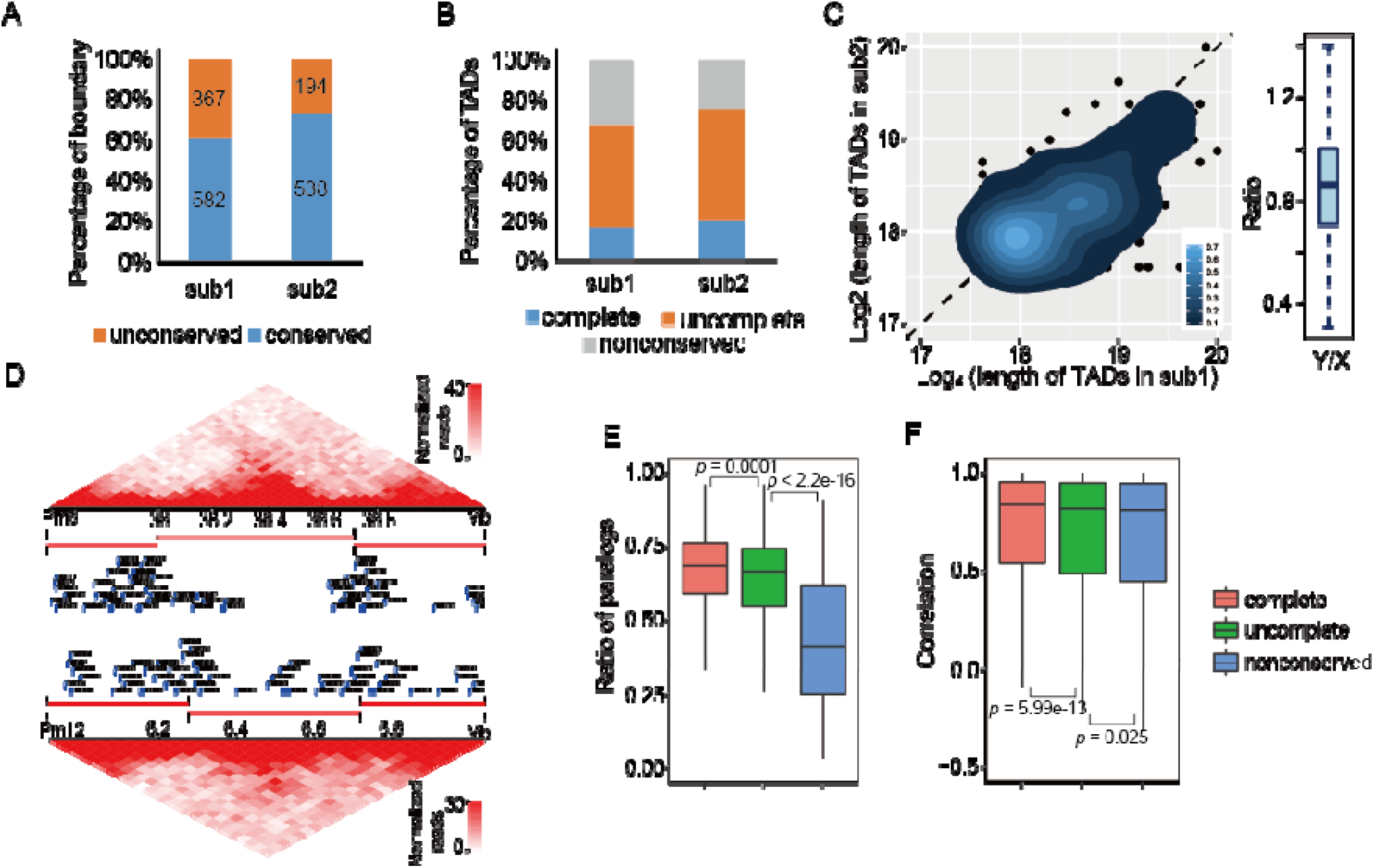
Identification of conserved TADs between two subgenomes of broomcorn millet, related to Fig2. **(A)** Percentages of conserved and non-conserved TAD boundaries between two subgenomes of broomcorn millet. Numbers on the bars indicated the absolute number of TAD boundaries. **(B)** The bar plot showed the proportion of non-conserved, incompletely conserved and completely conserved TADs between two subgenomes of foxtail millet. **(C)** The heatmap showed the lengths comparison of conserved TADs between two subgenomes of broomcorn millet. The box plot on the right of heat map showed the ratio of conserved TAD length between two subgenomes of broomcorn millet. **(D)** Example of conserved TADs on a syntenic region between two subgenomes of broomcorn millet. TADs and syntenic genes are indicated by red lines and blue rectangles, respectively. **(E)** Comparison of the percentages of paralogs in non-conserved, incompletely conserved and completely conserved TADs between two subgenomes of broomcorn millet. **(F)** Comparision of the correlation of the expression in non-conserved, incompletely conserved and completely conserved TADs between subgenomes of broomcorn millet. gene two

**Supplemental Figure 9.**
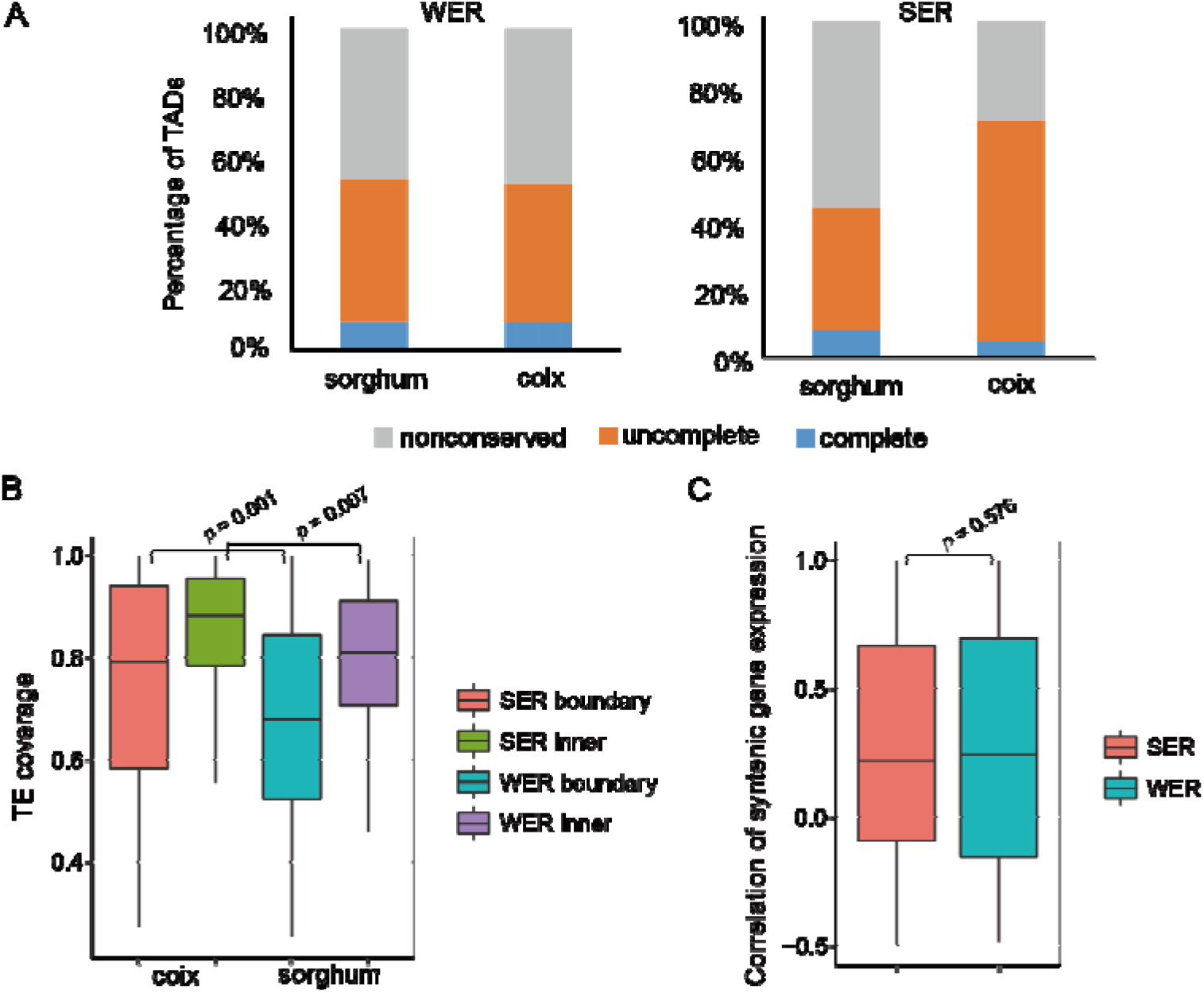
Charaterize the SER and WER between sorghum and coix, related to Fig3. **(A)** The bar plots showed the proportion of non-conserved, incompletely conserved and completely conserved TADs in SER and WER between sorghum and coix, with the corresponding numbers labeled on the bar. **(B)** Comparision of TE coverage of TAD boundary and inner region in SER and WER in coix. **(C)** Comparision of the correlation of gene expression in SER and WER between sorghum and coix.

**Supplemental Figure 10.**
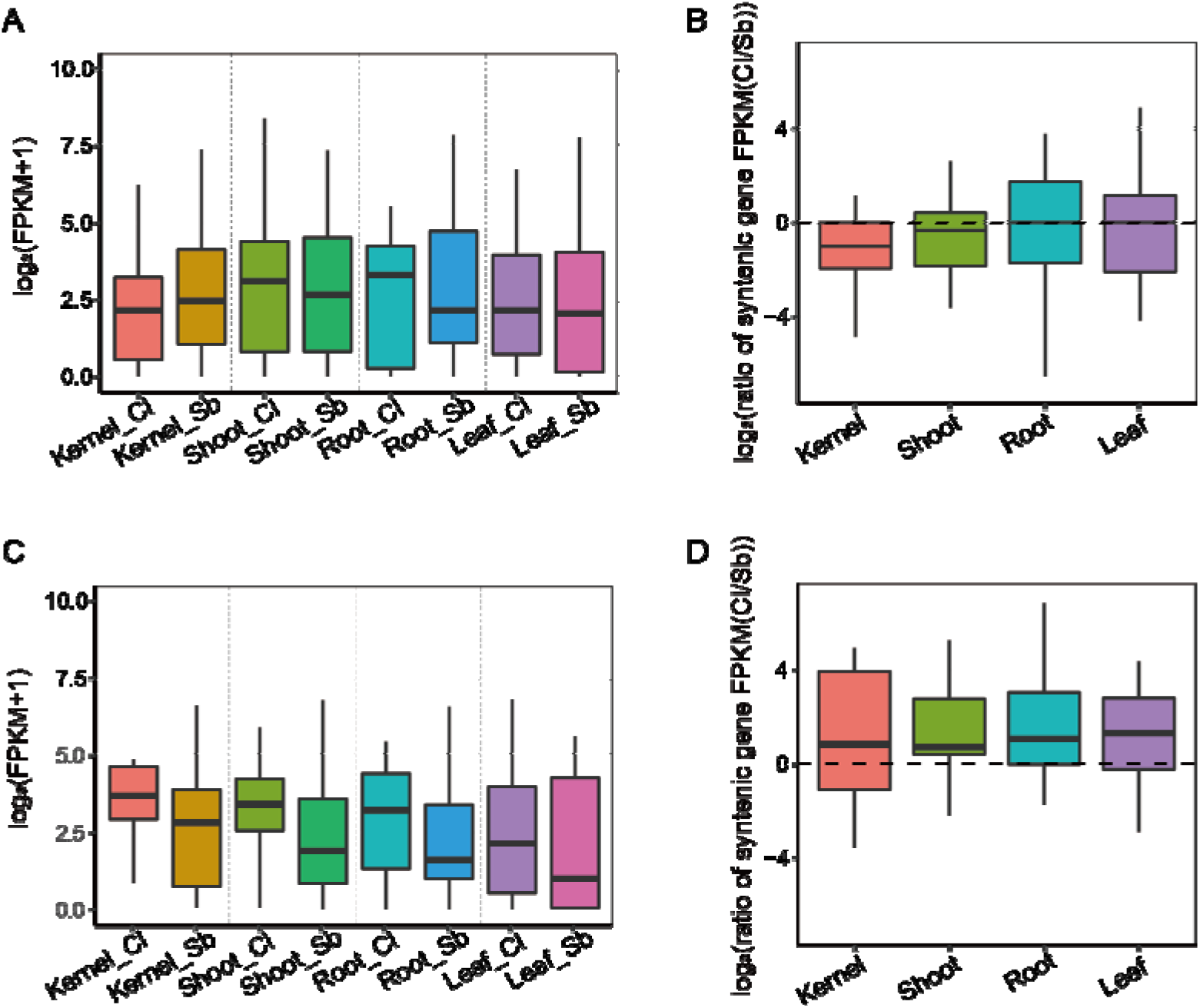
Comparision of the gene expression around the breakpoints of inversion regions between sorghum and coix, related to Fig3. (A, **B)** Comparison of the expression levels of genes in the TADs containing the breakpoints of inversion regions, which the breakpoints were located in inner region of TAD in Coix, and the corresponding breakpoints were in boundary region of TAD in Sorghum. **(C, D)** Comparison of the expression levels of genes in the TADs containing the breakpoints of inversion regions, which the breakpoints were located in boundary region of TAD in Coix, and the corresponding breakpoints were in inner region of TAD in Sorghum. The x-axis indicates different tissues and the y-axis indicates log_2_(FPKM+1) values in (A, C) and log2(Ratio of syntenic gene expression FPKM(Cl/Sb)) values in (B, D).

**Supplemental Figure 11.**
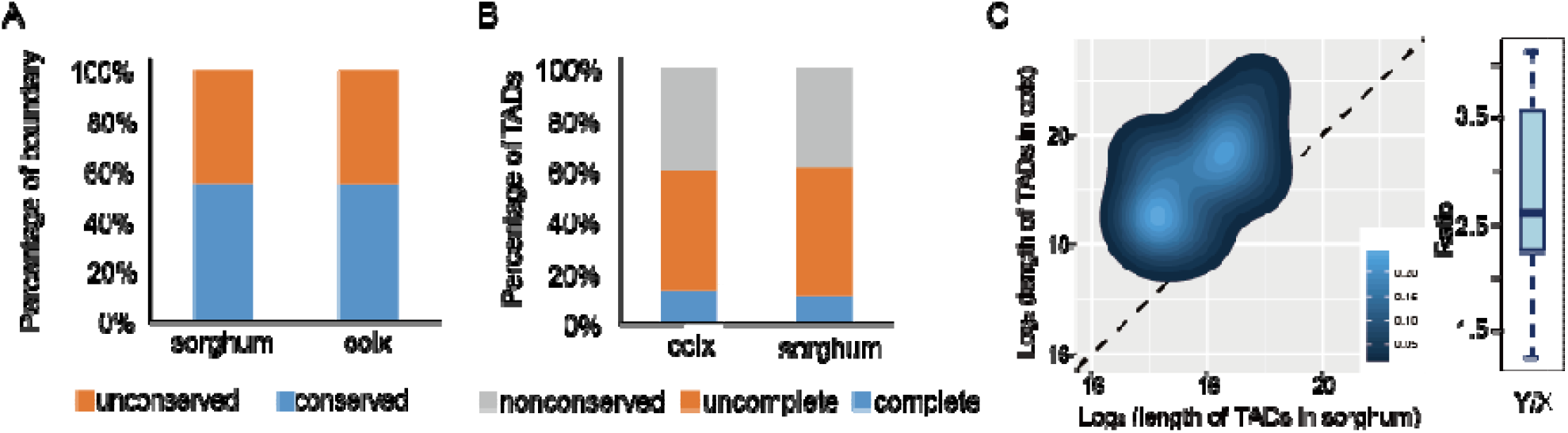
Identification of conserved TADs in the inversion regions between sorghum and coix, related to Fig4. **(A)** Percentages of conserved and non-conserved TAD boundaries in the inversion regions between sorghum and coix. Numbers on the bars indicated the absolute number of TAD boundaries. **(B)** The bar plot showed the proportion of non-conserved, incompletely conserved and completely conserved TADs in the inversion regions between sorghum and coix. **(C)** The heatmap showed the lengths comparison of conserved TADs in the inversion regions between sorghum and coix. The box plot on the right of heat map showed the ratio of conserved TAD length between coix and sorghum.

**Supplemental Figure 12.**
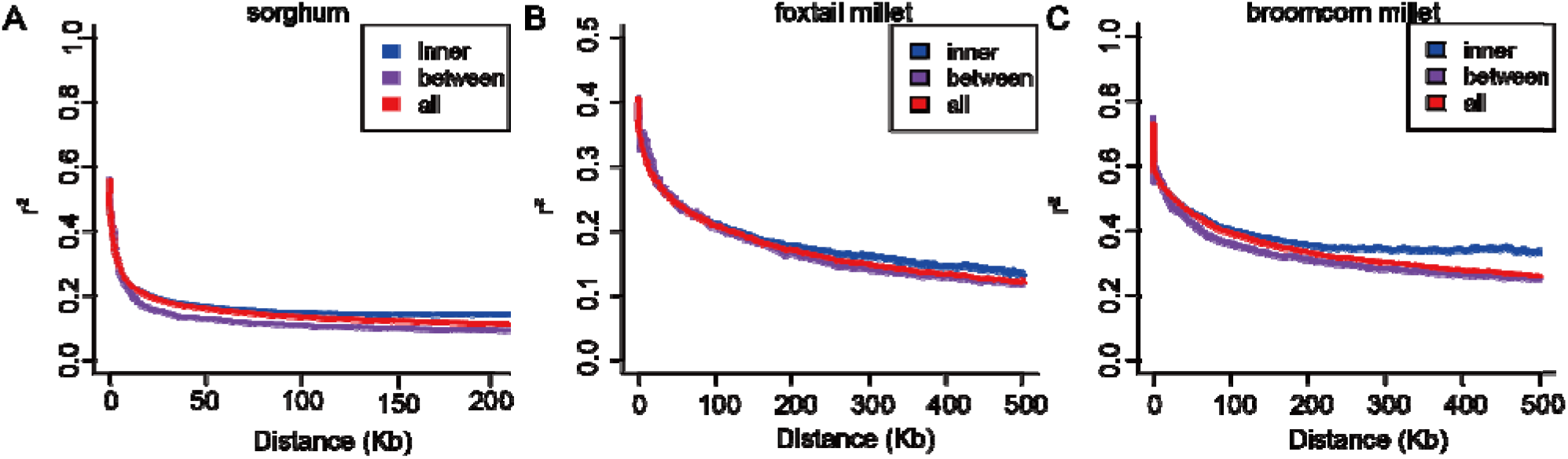
The LD decay between and within TADs in sorghum (A), foxtail mille (B) and broomcorn millet (C), related to Fig5.

**Supplemental Figure 13.**
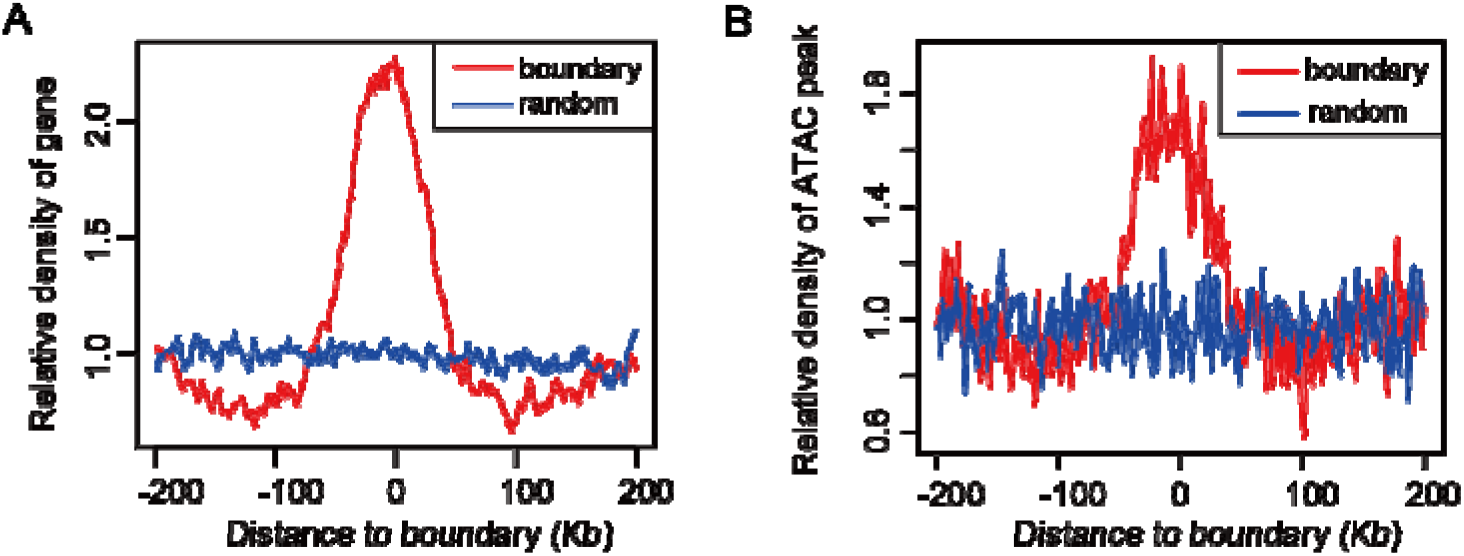
Gene and ATAC peak showed an enrichment at TAD boundary in maize, related to Fig5. **(A)** The relative density of gene around TAD boundary in maize. **(B)** The relative density of ATAC-seq peak around TAD boundary in maize.

**Supplemental Figure 14.**
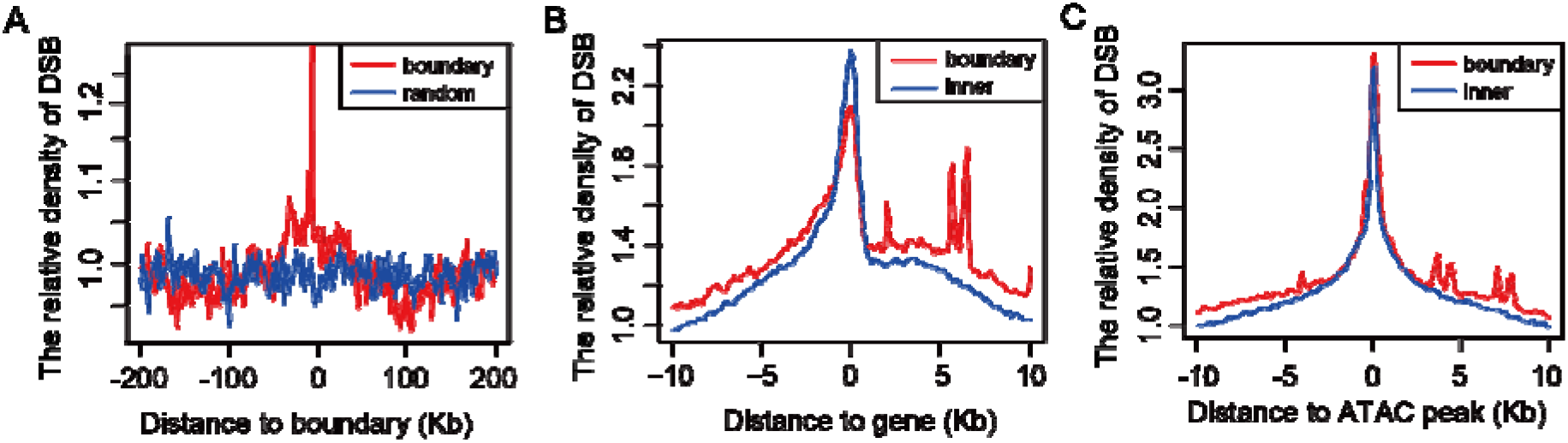
The relative distribution of DSB and SNP around TAD boundary and inner in maize, related to Fig5. **(A)** The relative density of DSB around TAD boundary in maize. **(B)** Compare the relative density of DSB around gene located in boundary and inner of TADs.**(C)** Compare the relative density of DSB around ATAC peak located in boundary and inner of TADs. **(D)** The relative density of SNPs around TAD boundary in maize.

**Supplemental Figure 15.**
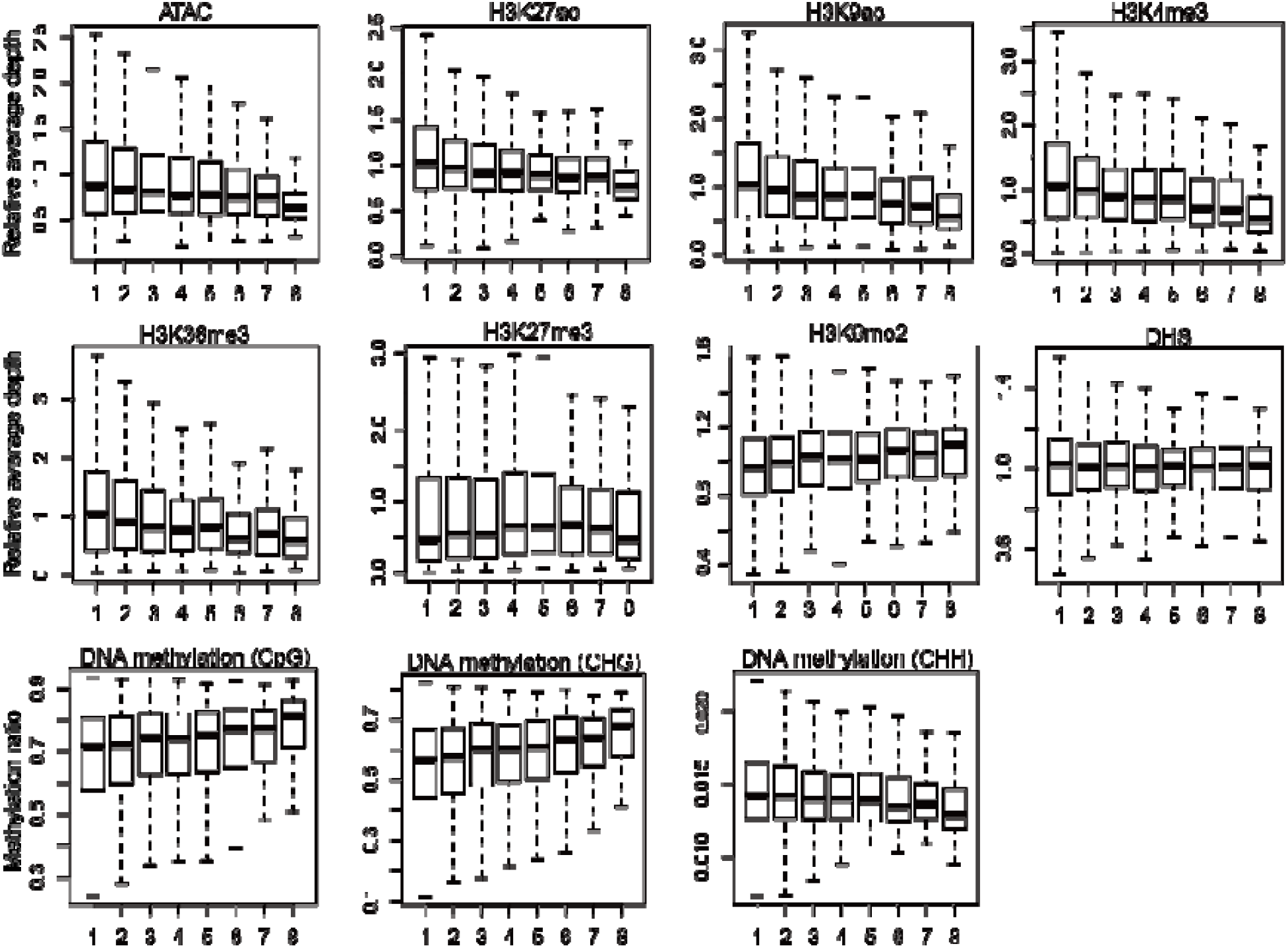
Comparison of various epigenetic features of chromatin located inside TADs with different sizes, related to Fig5. For each plot, the horizontal line indicates the mean value of the epigenetic feature across the genome. TADs in the boxplots were grouped according to their sizes as follow: group 1: TADs ≤ 300 Kb; group 2: 300 Kb < TADs ≤ 400 Kb; group 3: 400 Kb < TADs ≤ 500 Kb; group 4: 500 Kb <TADs ≤ 600 Kb; group5: 600 Kb < TADs ≤ 700 Kb; group 6: 700 Kb< TADs ≤ 800Kb; group 7: 800 Kb <TADs ≤ 1 Mb; group 8: TAD > 1 Mb.

**Supplemental table. 1.** Summary of TAD number and length at the corresponding resolution in five crop species.

|  | sorghum | coix | maize | foxtail millet | broomcorn millet |
| --- | --- | --- | --- | --- | --- |
| No. of chromosomes | 10*2 | 10*2 | 10*2 | 9*2 | 9*4 |
| Ploidy | diploid | diploid | diploid<br>(ancient<br>tetraploid) | diploid | tetraploid |
| Size of genome | 684Mb | 1.62Gb | 2.2Gb | 401Mb | 839Mb |
| No. of TADs | 1920 | 1965 | 4873 | 1679 | 3861 |
| Median TAD Length<br>(Kb) | 240 | 660 | 400 | 360 | 360 |

